# Single-molecule and super-resolved imaging deciphers membrane behaviour of onco-immunogenic CCR5

**DOI:** 10.1101/2022.05.19.492692

**Authors:** Patrick Hunter, Alex L. Payne-Dwyer, Michael Shaw, Nathalie Signoret, Mark C. Leake

**Affiliations:** Department of Biology, University of York, YO10 5DD, UK; School of Physics, Engineering and Technology, University of York, YO10 5DD, UK; National Physical Laboratory, Hampton Road, Teddington, Middlesex, TW11 0LW; Department of Computer Science, University College London, London, WC1E 6EA, UK; Hull York Medical School, University of York, YO10 5DD, UK

## Abstract

The ability of tumors to establish a pro-tumorigenic microenvironment is becoming an important point of investigation in the search for new therapeutics. Tumors form microenvironments in part by the ‘education’ of immune cells attracted via chemotactic axes such as that of CCR5-CCL5. Further, CCR5 upregulation by cancer cells, coupled with its association with pro-tumorigenic features such as drug-resistance and metastasis, has suggested CCR5 as a target for therapeutic inhibition. However, with several conformational “pools” being reported, phenotypic investigations must be capable of unveiling heterogeneity. Addressing this challenge, we performed structured illumination (SIM) and Partially TIRF coupled HILO (PaTCH) microscopy for super-resolution imaging and single-molecule imaging of CCR5 in fixed cells. Determining the positions of super-resolved CCR5 assemblies revealed a non-random spatial orientation. Further, intensity-tracking of assemblies revealed a distribution of molecular stoichiometries indicative of dimeric sub-units independent of CCL5 perturbation. These biophysical methods can provide important insights into the structure and function of onco-immunogenic receptors and a plethora of other biomolecules.

**Highlights:**

- We use SIM and novel PaTCH microscopy for precise bioimaging and single-molecule tracking of receptor protein CCR5 in model cell line
- By tracking the position of CCR5 assemblies we conclude that they are clustered in the plasma membrane beyond a level expected from a random distribution
- We use these high-precision data to determine molecular stoichiometries of CCR5 assemblies

## Introduction

The mechanism by which cancer cells obtain immune evasive features remains an open question. However, current research indicates that cancer cells are able to manipulate chemokine networks to support tumor progression, with the main chemotactic axis utilised being that of C-C chemokine receptor type 5 (CCR5) and its ligand C-C chemokine ligand type 5 (CCL5) (Aldinucci et al., 2020; Jiao et al., 2019; Upadhyaya et al., 2020). CCR5 is a member of the seven transmembrane family of G protein coupled receptors (GPCRs) and is found in various white blood cells including T-cells and macrophages. CCR5 responds to a range of chemokines, such as CCL5, known to induce chemotaxis towards sites of immune response (Aldinucci and Colombatti, 2014). Despite this immunogenic feature of the CCR5-CCL5 axis, cancer cells are capable of “educating” migrating immune cells to form an immunosuppressive tumor microenvironment (Aldinucci et al., 2019; Chang et al., 2012; Tan et al., 2009). Further, oncogenic transformation of cells has been shown to increase the surface expression of CCR5 (Erreni et al., 2009; Mañes et al., 2003; Sales et al., 2014; Vaday et al., 2006). As a result, CCR5 has become a strong point of investigation in studies relating to both immunology and cancer.

Studies have linked the upregulation of CCR5 by cancer cells with poor prognosis for patients, with the CCR5-CCL5 axis being found to aid in tumor growth, metastasis and drug-resistance amongst other pro-tumorigenic features (Aldinucci et al., 2008; Aldinucci and Casagrande, 2018; Vaday et al., 2006). These investigations have culminated in the consideration of the CCR5-targeted drug Maraviroc, previously utilised as a therapeutic for HIV, to be repurposed as a clinical treatment for cancer patients (Aldinucci et al., 2020; Haag et al., 2020; Miao et al., 2020). It is clear that the CCR5-CCL5 axis plays an important role in both immune dynamics and tumor progression, and that investigations into the behavior of CCR5 along the surface membrane are likely to be beneficial in the development of new therapeutics.

Previous studies investigating the antigenic behaviour of CCR5 suggest the existence of multiple conformational subpopulations of CCR5, an observation that is derived from the characteristic recognition of structurally distinct ligands by varying proportions of cell surface CCR5 (Colin et al., 2018, 2013; Fox et al., 2015; Weichseldorfer et al., 2022). Further, CCR5 has been shown to exist in both homodimeric and heterodimeric states, with the homodimerization of CCR5 being shown to take place in the endoplasmic reticulum before reaching the cell surface (Jin et al., 2018), any downstream clustering would therefore be expected to carry a periodicity in apparent stoichiometry of two molecules (Martínez-Muñoz et al., 2018b). Considering that the various purported conformational and oligomeric states of CCR5 exhibit downstream effects on the receptor chemotactic functionality (Berro et al., 2011; Colin et al., 2013), the nature of this heterogeneity requires further investigation.

Although earlier single-molecule studies have been carried out using GPCRs (Joseph et al., 2021; Kasai and Kusumi, 2014; Tian et al., 2017), previous investigations into the oligomeric status of CCR5 in particular have been focused within the bulk ensemble regime, utilising techniques such as standard fluorescence confocal microscopy, flow cytometry and Western blotting (Colin et al., 2018; Jin et al., 2018). Through the extension of cell-surface CCR5 investigations to include 3D super-resolution microscopy and high-speed single-molecule tracking, here we have been able to visualise CCR5 expression with a spatial resolution twice that of the optical diffraction limit as well as provide stoichiometry estimates based on single-molecule measurements, thereby unveiling the heterogeneity lost by bulk ensemble techniques. Further, studying the effects of ligand perturbations make it possible to draw additional conclusions regarding the effects of CCL5 binding on the cell surface expression of CCR5.

Here, we introduce a study which utilises the accompaniment of two light microscopy techniques, chosen for their high spatial and temporal resolution, to study Chinese hamster ovary (CHO) cell lines which have undergone genetic modification for the production of CCR5 in the absence of endogenously produced CCR5. Our study employs structured illumination microscopy (SIM) for the investigation of cell surface CCR5 distributions within CHO-CCR5 cells, a cell line utilised in multiple previous studies (Mack et al., 1998; Signoret et al., 2000, 2004, 2005). Also, through the creation of a new line of transfected CHO cells, which express CCR5 N-terminally fused to GFP to a level compatible with single-molecule microscopy, it has been possible to provide single-molecule stoichiometry estimates using a newly developed microscopy technique that combines the high signal-to-noise ratio of total internal reflection fluorescence (TIRF) microscopy whilst benefiting from the increased penetration depth of Highly Inclined and Laminated Optical sheet (HILO) microscopy (Tokunaga et al., 2008). This new imaging mode, that we denote Partially TIRF coupled HILO (PaTCH) microscopy is distinguished from TIRF and HILO through the employment of an optimised angle of incidence which aims to improve the study of transmembrane proteins in cells exhibiting a complex basal membrane topology. TIRF relies on an angle of incidence which surpasses the critical angle of a glass-water interface, ensuring total internal reflection of the excitation beam at the coverslip and thereby resulting in an exponentially decaying evanescent field that extends a short distance through the sample. This evanescent field allows for the signal enhancement of molecules only within a few hundred nanometers of the coverslip surface whilst reducing intracellular background. Conversely, HILO employs a lower angle of incidence which results in the transmission of the excitation beam through the sample at a specified angle, reducing the excitation of out-of-focus layers and back-scattered light, and is therefore typically used for intracellular imaging with lower background albeit without benefiting from any signal enhancement. By employing an intermediate angle, PaTCH aims to produce an excitation field that is coupled jointly into TIRF and HILO modes, thereby providing an increased signal from molecules within the basal membrane when compared with HILO whilst allowing for the excitation of molecules above the limited excitation field achieved by TIRF (see Supplementary Figure 1). Finally, using this new technique we then demonstrate how these methods can be used to investigate the surface expression and aggregation of CCR5 after exposure to the receptor’s CCL5 ligand.’

## Results

### SIM reveals CCR5 as distinct puncta distributed throughout the cell membrane in 3D

Initial investigations aimed to visualise the distribution of CCR5 in CHO-CCR5 cells, for which previous bulk ensemble studies had been performed. For this purpose, SIM was used to acquire super-resolution images of CHO-CCR5 labeled with the CCR5 mAb MC-5, itself labeled with DyLight 650 (Figure 1 a-g). Representation of these images included the correction of photobleaching effects on the fluorescence intensity, the exclusion of background fluorescence located outside the cell, as well as the inclusion of magnified insets of the CCR5 puncta. From these images we see that membrane-bound CCR5 assemblies appear as distinct puncta throughout the entire plasma membrane, with puncta appearing uniformly distributed along the basal membrane. Further, due to the external labelling of CCR5, in cross-sectional images corresponding to planes above the basal membrane puncta are only visible within annulus-shaped regions of the cell surface.

**Figure 1.**
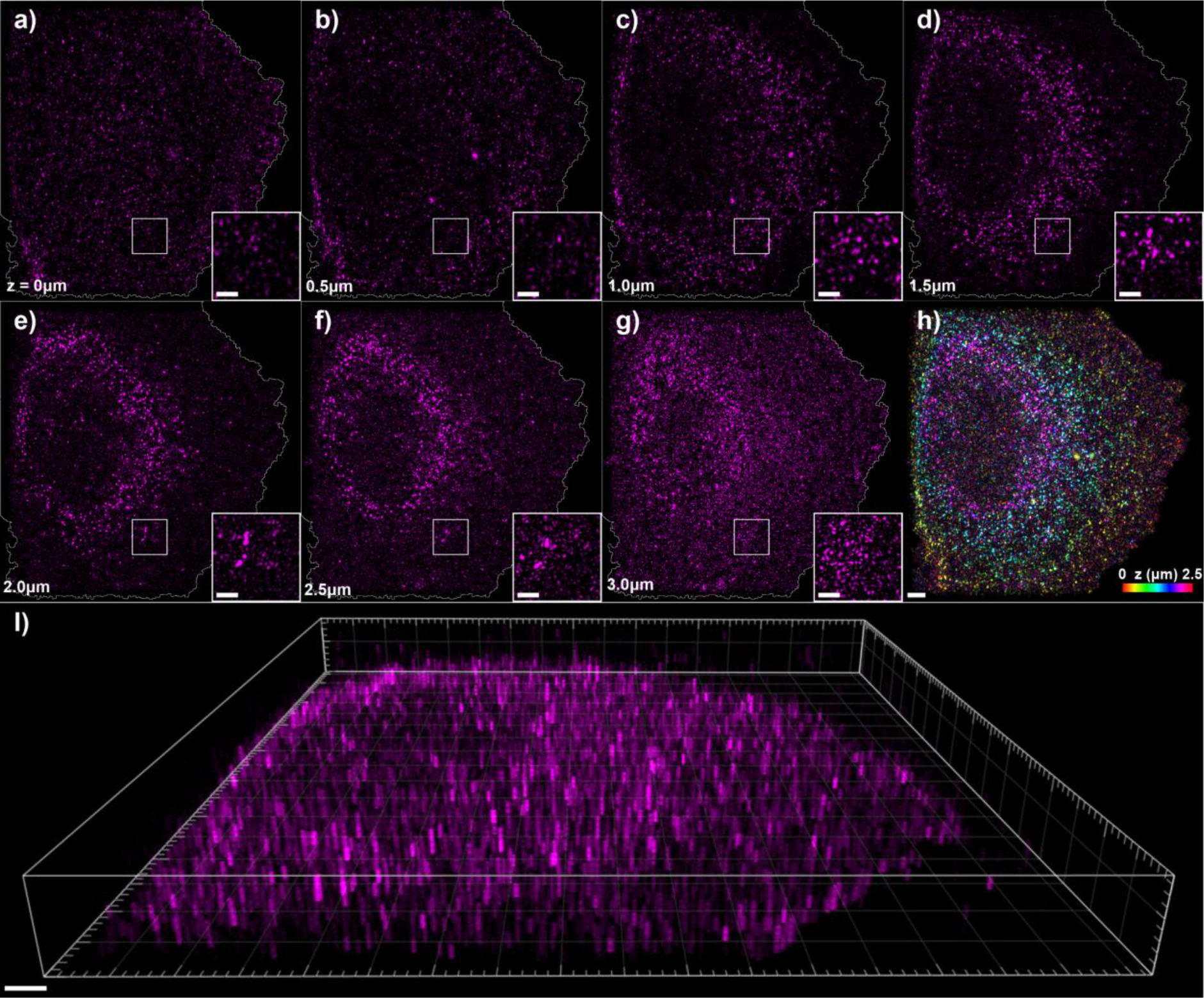
SIM images of CHO-CCR5 cells labeled with DyLight 650-MC-5. *a - g) Individual SIM image planes of a CHO-CCR5 cell, from the basal membrane through to the apical membrane in 500 nm steps, displaying cell boundary segmentation and magnified insets. h) Color depth projection over the images shown in a-f). i) 3D reconstruction of cell images shown in a-g). (Scale bar 2 μm, (magnified insets 1 μm))*.

Despite the increase in background fluorescence seen in Figure 1 g) due to photobleaching correction, the annular distribution of fluorescent foci is clearly retained. Finally, acquisition of optically sectioned slices allowed the reconstruction of color depth projections, as well as 3D images and movies, as shown in Figure 1 h, i) and Supplementary Movie 1. These reconstructions enable the visualisation of this annular distribution of cell surface CCR5, thereby revealing the topology of the cellular exterior while further highlighting the uniformity of the CCR5 distribution.

### Clustering behavior of CCR5 assemblies is revealed through Ripley’s H function-based analysis

Mammalian cells present a variety of cell morphologies, including membrane protrusions which can be caused by a stress response to sample preparation. Although these filopodia are not of high biological relevance to this investigation, the extensions of the representative cell shown in Figure 2 a-d) provide a clear illustration of the increased image quality (higher spatial resolution and image contrast) of SIM over traditional widefield microscopy. This improved image quality, coupled with the optical sectioning property of SIM, aid quantitative analysis of image data and enable puncta of CCR5 assemblies to be isolated from the cellular background through intensity thresholding, as shown in Figure 2 c). Closer investigation of these isolated puncta reveals a spatial distribution with distinct ‘hot spots’ as shown in Figure 2 d). These results indicate the existence of spatial clustering of individual puncta. To analyse this further, we performed clustering analysis using Ripley’s *H* function (Kiskowski et al., 2009) on the centroids of isolated clusters. As represented in Figure 2 e), this method is based on the counting of objects at increasing distance averaged over all possible origin points inside the cell, and can be used to describe the level of clustering, uniformity or dispersion of points. By visualising the Ripley’s *H* values as a function of radius spanning the diameter of the cell’s basal membrane, we see that net clustering of points is characterised by accumulation of positive Ripley’s *H* values across the cell, as shown in Figure 2 f-g). The modal clustering gradient thereby provides an estimation of the spatial correlation of CCR5 puncta throughout the cell, with the modal average gradient of CHO-CCR5 cells found to be positive: *dH(r)/dr* = 0.004 ± 0.001. The initial minima displayed in the Ripley’s *H* value distribution provides an estimate for the nearest neighbor distance between CCR5 puncta, with the average for CHO-CCR5 cells measured at 290 ± 10 nm.

**Figure 2.**
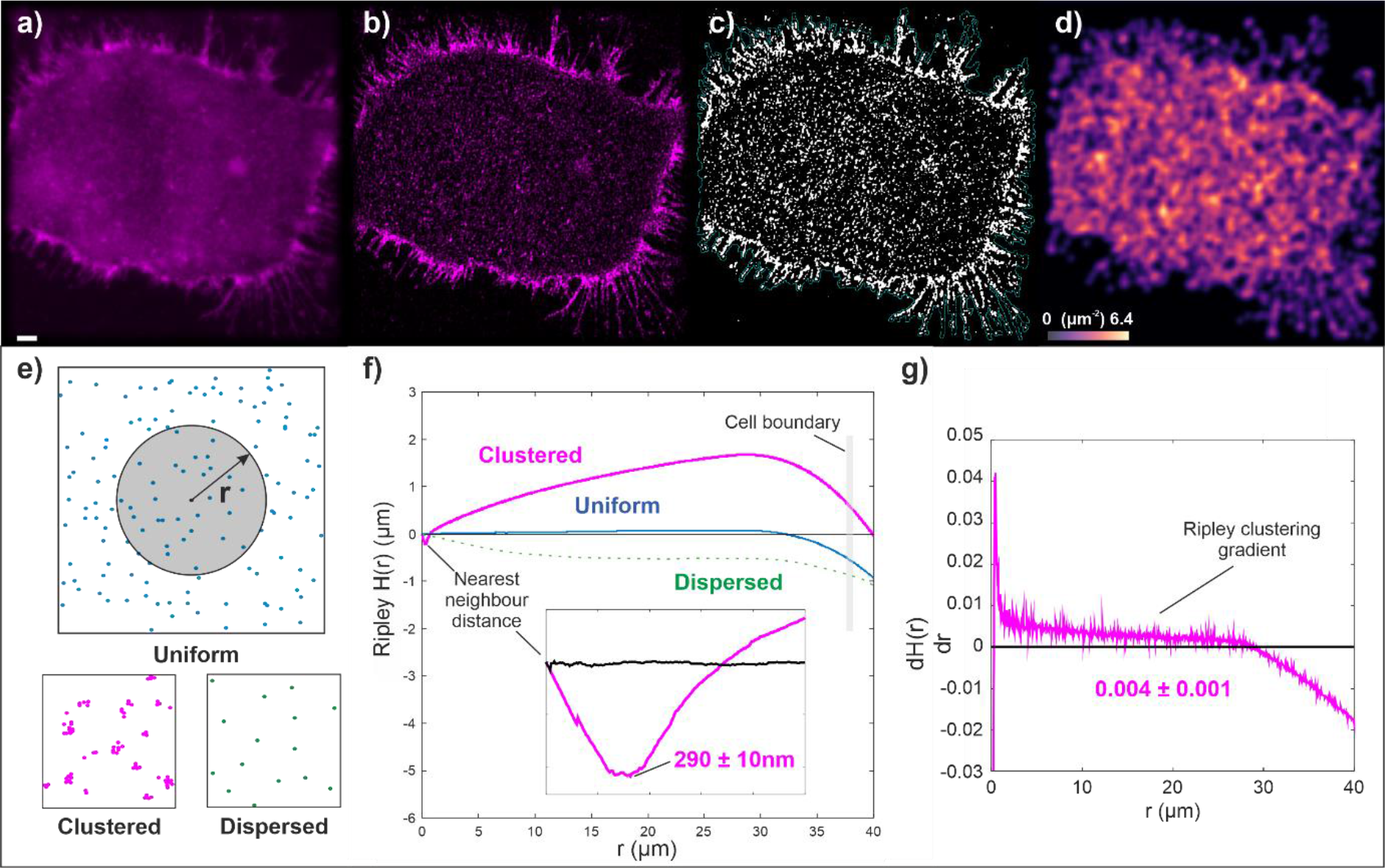
Images and quantitative clustering metrics of CCR5 in the basal membrane of CHO-CCR5 cells labeled with DyLight 650-MC-5. *a) Widefield image of the basal membrane of a representative CHO-CCR5 cell (scale bar 2 μm). b) SIM reconstruction of the same cell. c) Binarized mask of the SIM image displaying thresholded CCR5 puncta. d) Number density distribution of individual puncta centroids, ranging from 0 - 6.4 μm^−2^. e) Illustration demonstrating the function of Ripley’s H analysis in the investigation of uniform, clustered and dispersed point distributions. f) Distribution of Ripley’s H values for a real representative CHO-CCR5 cell (magenta) and a generated 2D map of points whose displacement separations are sampled from a random Poisson distribution (blue). Also shown is an illustrative curve depicting the expected results of a generic dispersed point distribution (green dotted). Inset shows the same distribution of Ripley’s H values for the CHO-CCR5 cell (magenta) and random point distribution (blue) over a range of 0 - 1 μm and provides an average nearest-neighbor distance for binarized CHO-CCR5 puncta (N=10 cells). g) The gradient of the distribution of Ripley’s H values for the CHO-CCR5 cell represented in f) which provides a modal average clustering gradient for CHO-CCR5 (N=10 cells)*.

### Clones of GFP-CCR5 CHO cells developed for PaTCH microscopy and characterised using flow cytometry

To determine structural characteristics of the CCR5 assemblies, we performed single-molecule PaTCH microscopy-based studies to measure the number of molecules present in assemblies, which we denote as the stoichiometry. While PaTCH benefits from the simplicity of constitutively fluorescent probes and does not require photoswitchable or photoactivatable probes (Jin et al., 2021; Yuan et al., 2021), the successful investigation of individual CCR5 assemblies using PaTCH microscopy does rely on fluorescent proteins being expressed to a level compatible with single-molecule localization microscopy. For this purpose, CHO cells transfected to express GFP-CCR5 fusion proteins underwent single-cell cloning to create several populations with varying levels of expression. Figure 3 shows one such population, optimised for single-molecule imaging. In Figure 3 a-c) we see that despite exhibiting low expression suitable for single-molecule studies, the GFP expression of the positive sample is significantly higher than that of the control in both flow cytometry and microscopy-based experiments, providing confidence in this new model for CCR5 investigations. Additionally, to confirm the functionality of CCR5 in this newly created cell line, we performed a calcium flux assay (see Supplementary Figure 2 a). These results reveal CCL5-induced calcium mobilisation in agreement with previous studies (Gómez-Moutón et al., 2004), albeit at a level that is representative of the lower CCR5 expression exhibited by this cell line.

**Figure 3.**
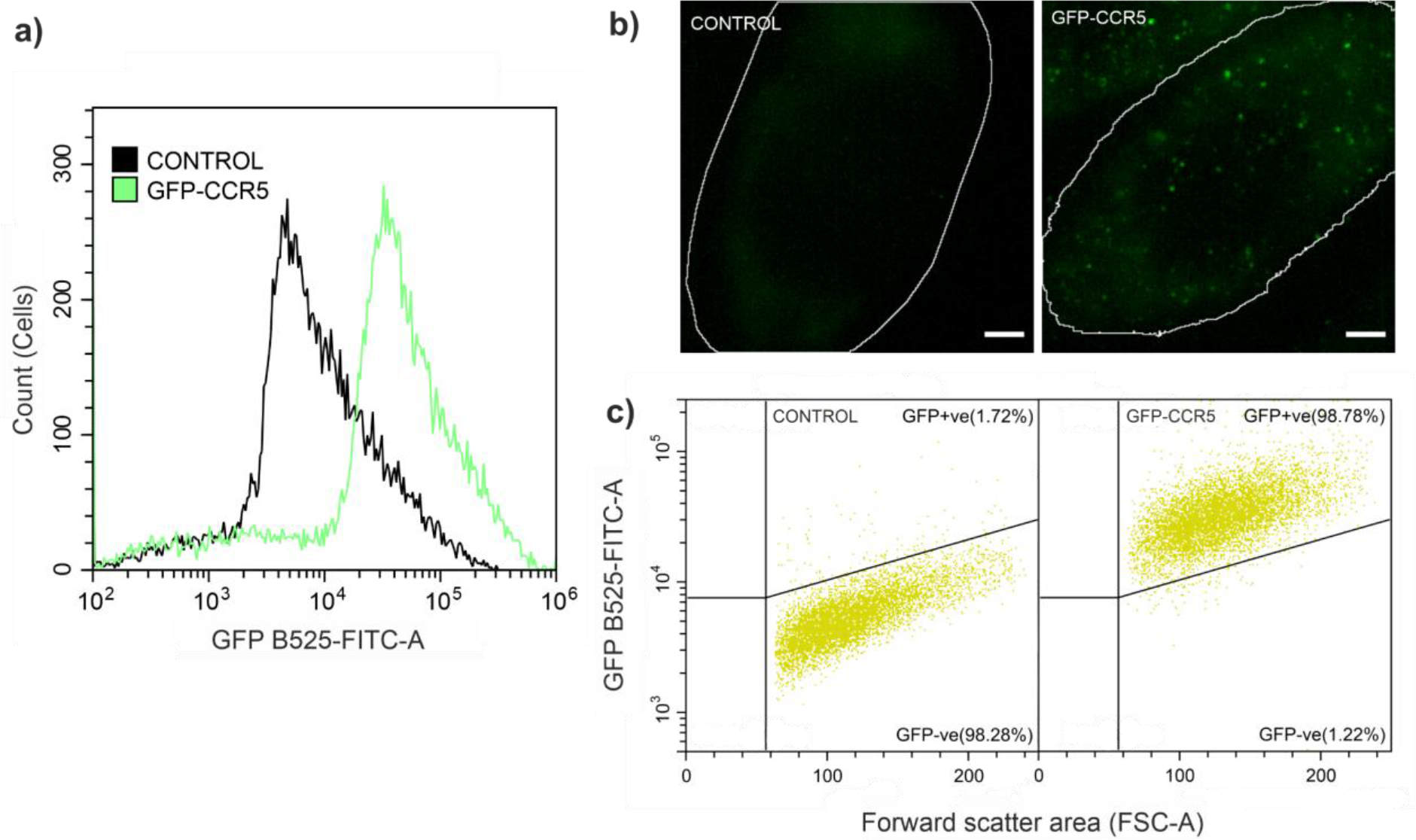
Characterisation of GFP-CCR5 expression in transfected CHO cells using flow cytometry and PaTCH microscopy confirms increase above control samples. *a) Distribution of GFP emission intensity over several hundred cells for both control and GFP-CCR5 positive samples. b) PaTCH images of the basal membrane of both control and GFP-CCR5 positive cells. Cell boundary segmentation shown in white. (Scale bar 2 μm). c) Scatterplot of GFP-range emission intensity against the forward scatter area of both control and GFP-CCR5 positive cells. Both a) and c) show cell populations after gating to remove debris and doublets*.

### PaTCH investigation of basal membrane GFP-CCR5 also reveals CCR5 assemblies as small puncta

As can be seen in Figure 4 a), despite a reduction in expression, GFP-CCR5 forms distinct puncta across the basal membrane of transfected CHO cells in a qualitatively similar distribution to that of DyLight 650-MC-5 labeled CCR5 in CHO-CCR5 cells, as shown in Figure 4 b). Althoug the increased spatial resolution of SIM images provides improved segmentation of CCR5 puncta for the determination of spatial clustering, high-speed PaTCH microscopy allows the determination of time-dependent processes such as dye photobleaching effects. Although further studies are required into the accurate determination of puncta diameter in both respective cell models, the general uniformity in the size of CCR5 puncta between both cell models provides confidence that any spatial dependence in CCR5 expression between these two models is related.

**Figure 4.**
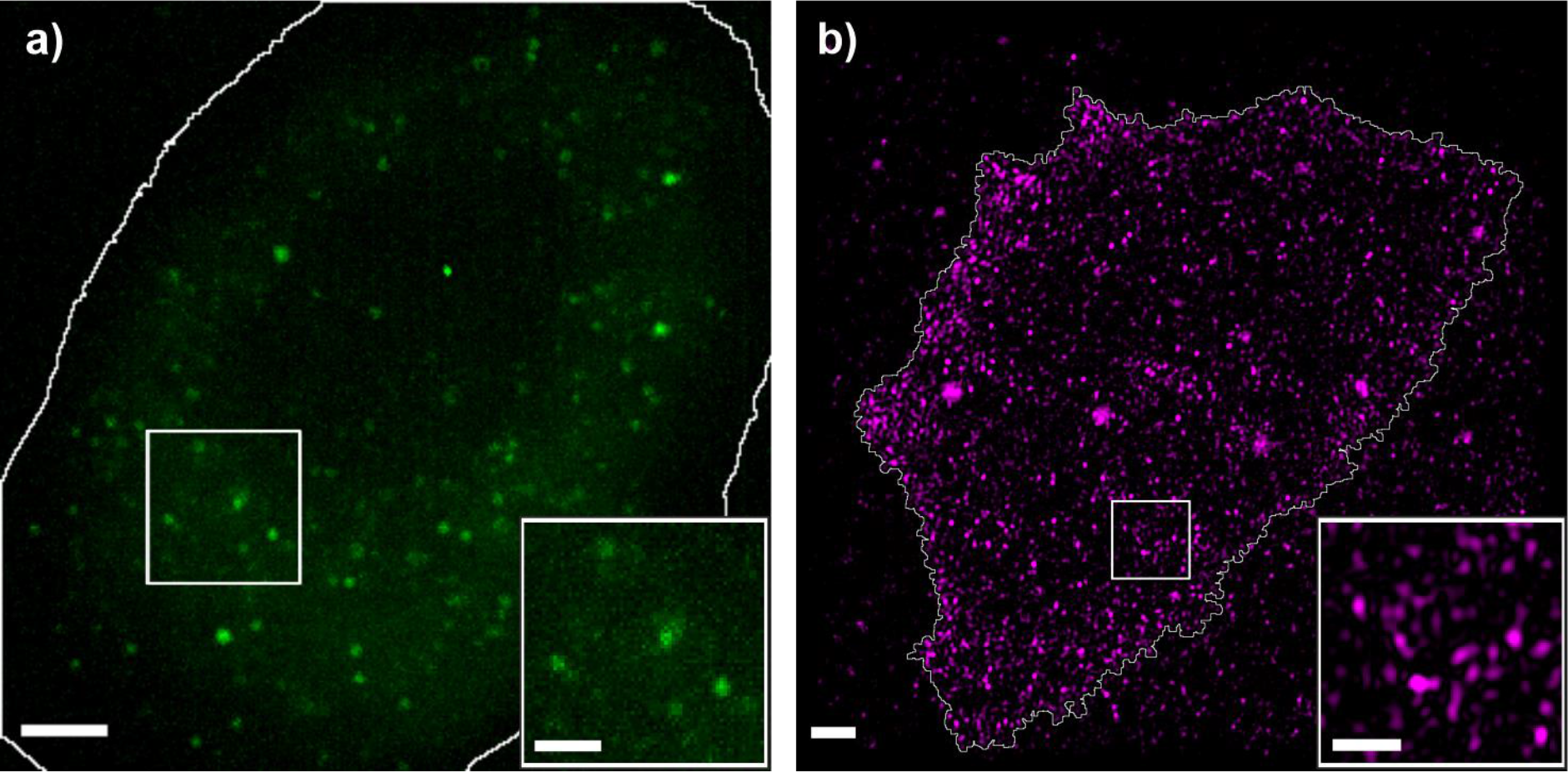
Comparison of GFP-CCR5 CHO cells with established CHO-CCR5 cells. *Fluorescent labels shown to be distributed in distinct puncta across cellular membranes in a) GFP-CCR5 expressing CHO cells imaged using PaTCH microscopy and b) DyLight 650 labeled CHO-CCR5 cells imaged using SIM. Cell boundary segmentation shown in white. (Scale bar 2 μm, (magnified insets 1 μm))*.

### Stoichiometry measurements reveal CCR5 assemblies to comprise homodimers

Utilising the single-molecule sensitivity of PaTCH microscopy, we are able to identify the characteristic brightness of a single GFP as the modal intensity of single GFP-CCR5 molecules following sufficient photobleaching, further confirmed using recombinant GFP and the quantification of single-molecule photobleaching steps in fluorescence intensity (see Supplementary Figure 3 b). This global value is used to normalise the initial intensity of GFP-CCR5 foci in order to acquire estimates of stoichiometry. A wide distribution of stoichiometries is revealed by collating all detected tracks across all GFP-CCR5 CHO cells (represented as a kernel density estimate (Leake, 2014) in Figure 5). This population of independent track-derived stoichiometries shows characteristic peaks, with the average nearest-neighbor interval between independent stoichiometry measurements revealing the typical periodicity inside oligomeric assemblies, if such periodicity exists (Wollman et al., 2017). Although the accurate measurement of small differences between two large stoichiometries is difficult, it is possible to successfully average those differences over many pairs (see Methods: Single particle tracking). Thereby, for CCR5 in the absence of ligand, we find the average periodicity to be 2.3 ± 0.5 CCR5 molecules (Figure 5 inset), indicating a strong tendency for CCR5 molecules to occur in dimeric subunits inside CCR5 assemblies.

**Figure 5.**
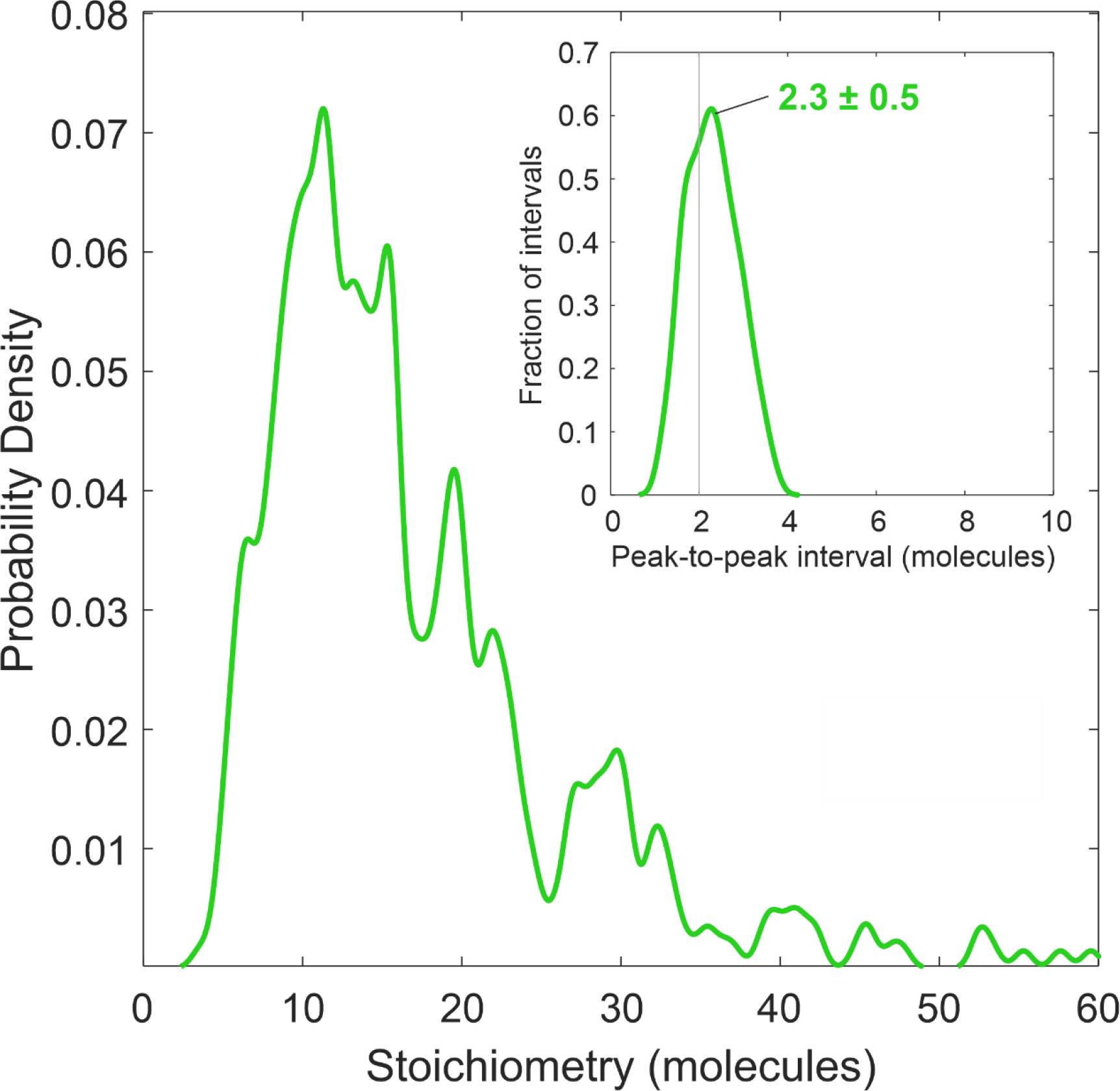
Periodic stoichiometry distribution indicates that CCR5 assemblies comprise dimeric subunits. *Kernel density estimates of stoichiometry and (inset) periodic stoichiometry intervals of GFP-CCR5 associated foci (N=460 tracks) detected by PaTCH microscopy in GFP-CCR5 transfected CHO cells (N=9 cells). Kernel width = 0.6 molecules, corresponding to the total uncertainty in the single molecule stoichiometry, rather than statistical fluctuations. Measured intervals in probability density or stoichiometry are therefore more reliable at lower stoichiometry*.

### Tracking of single molecules in PaTCH after addition of CCL5 indicates a broader spread of CCR5 foci stoichiometries

This study was extended by applying the above methods to study CCR5 upon stimulation with the agonist CCL5 at a concentration of 100 nM. CHO-CCR5 cells treated for 5 minutes with CCL5 were imaged using SIM microscopy as shown in Figure 6 a-i). As in Figure 1, representation of these images included the correction of photobleaching, the exclusion of background fluorescence located outside of the cell and the inclusion of magnified insets of the CCR5 puncta. However, this cell is displayed using slightly differing contrast settings to enable optimal image presentation, despite small qualitative differences in brightness between cells (see Supplementary Figure 4). In a similar manner to Figure 1, we see the expression of CCR5 assemblies as small puncta throughout the cell membrane, with CCR5 uniformly distributed across the basal membrane and visible within approximately annulus-shaped regions for imaging planes above the basal membrane, as visualised in 3D (see Supplementary Movie 2).

**Figure 6.**
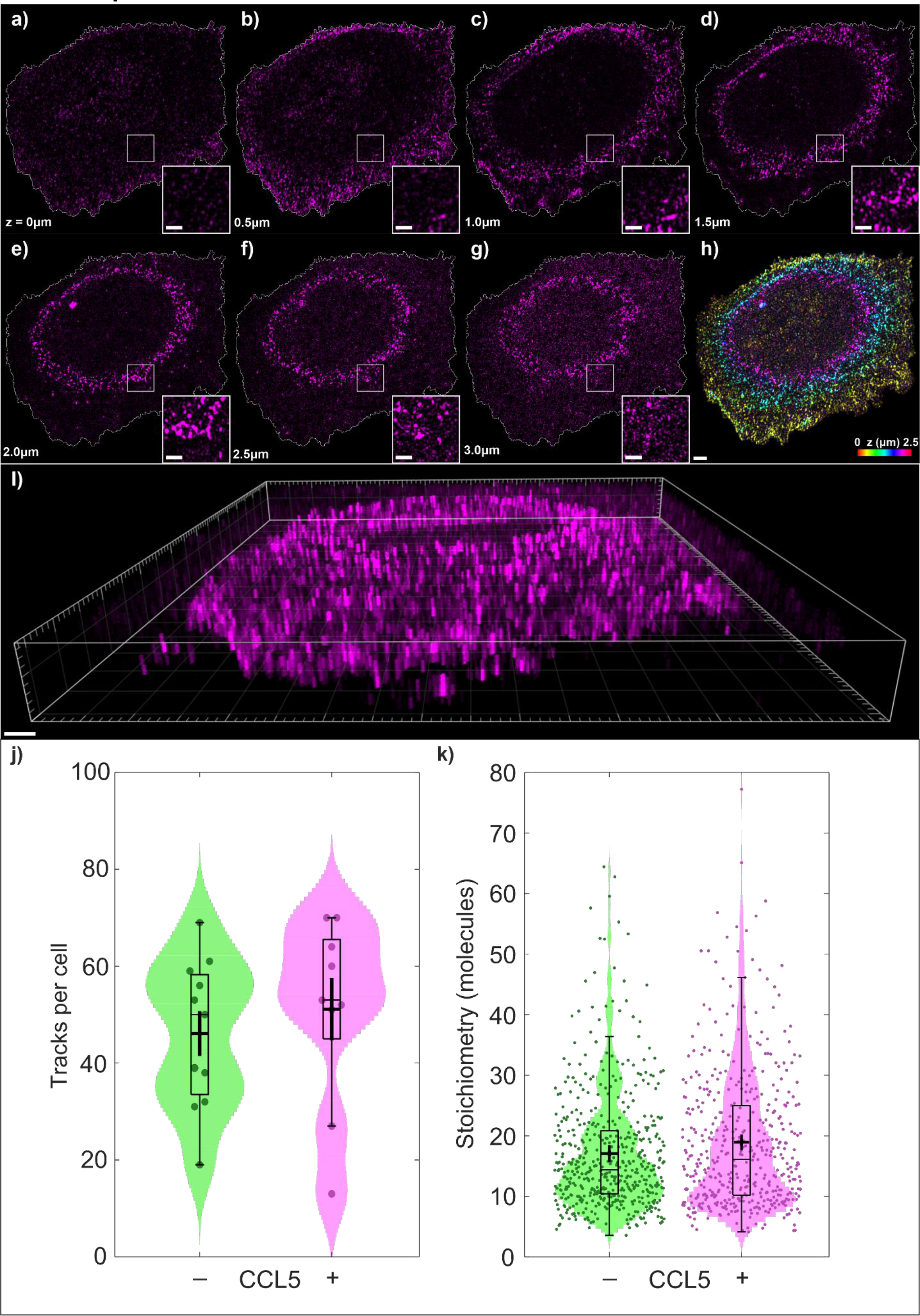
Investigating the perturbation of CCR5 with agonist CCL5 using SIM and PaTCH microscopy. *a - g) Individual SIM images of a CCL5 perturbed CHO-CCR5 cell, optically sectioned from the basal membrane through to the apical membrane in 500 nm increments, including cell boundary segmentation and magnified insets. h) Color depth projection of cell images shown in a -f). i) 3D reconstruction of cell images shown in a - g) (Scale bar 2 μm, (magnified insets 1 μm)). j) Numbers of GFP-CCR5 tracks detected per cell, with and without CCL5 perturbation, represented as dots spread horizontally to allow individual visualisation (N=9 or 11 cells, respectively). k) Violin distributions of the stoichiometry of GFP-CCR5 assemblies, with and without CCL5 perturbation, represented as dots spread horizontally to allow individual visualisation (N=460 or 507 tracks respectively). Bars, boxes and whiskers denote median, interquartile range (IQR) and 2.5× IQR respectively and the cross denotes the mean ± sem for both j and k*.

The interaction between CCR5 and CCL5 is a process reported to induce internalisation, with CCL5 perturbed cells being expected to exhibit a reduction in cell surface CCR5. Despite this, the SIM image data do not indicate any reduction in the overall number of puncta in perturbed versus non-perturbed cells. The PaTCH images provide quantitative detail which confirms that addition of CCL5 does not tend to change the average number of CCR5 assemblies, as the mean number of tracks we detect are not significantly different before and after CCL5 (46.1 ± 4.8 and 51.1 ± 6.8 tracks per cell respectively) under the Brunner-Munzel (BM) test (n=18, p=0.429 | not significant at adjusted p<0.01 level, NS). The mean stoichiometry of these assemblies lies in the vicinity of ~20 molecules regardless of CCL5 addition (BM test: 17.1 ± 0.4 and 18.6 ± 0.5 molecules before and after CCL5 addition respectively, n=920, p=0.145 |NS). Taken together, these results suggest the total amount of CCR5 presented on the cell surface is approximately conserved under our experimental conditions.

However, CCL5 appears to affect large and small assemblies of CCR5 differently. We see a larger spread of stoichiometry of assemblies in perturbed cells (Figure 6 j-k) than can be accounted for by any difference in sampling variance. At the lower end, assemblies are more commonly comprised of approximately 8 CCR5 molecules, while at the higher end CCR5 contributes towards the growth of larger assemblies of greater than approximately 36 molecules. These two sub-groups are populated at the expense of intermediate assemblies near the mean stoichiometry. This is confirmed by the difference in the means of each, before and after CCL5 addition (BM test at stoichiometry < 15 molecules, n=436, p=0.0085|*; at stoichiometry > 15 molecules, n=480, p=0.0053|*), despite almost identical numbers of assemblies in each group.

By studying CCL5 perturbation in GFP-CCR5 CHO cells using PaTCH microscopy, we determined an average periodicity in stoichiometry of 2.2 ± 0.3 CCR5 molecules in the assemblies after perturbation. This is consistent with the result prior to ligand exposure (Figure 5 inset), and strongly indicates that the consistent dimeric composition of CCR5 within assemblies is unaffected by CCL5 (see Supplementary Figure 5). Additionally, the effect of extended ligand perturbation was investigated using a potent analog of CCL5 known as PSC-RANTES (Escola et al., 2010; Fox et al., 2015) at a concentration of 100 nM (see Supplementary Figure 2 b). This flow cytometry-based assay monitored the accessibility of the chemokine binding site and the GFP moiety of GFP-CCR5 under varying duration of ligand exposure. This investigation revealed a reduction in both the accessibility of GFP and the epitope overlapping with the chemokine binding site, indicating ligand-binding and subsequent internalisation of GFP-CCR5. However, when compared with previous studies performed on CHO cells expressing wild-type CCR5 (Signoret et al., 2004), GFP-CCR5 exhibits a slower internalisation response.

## Discussion

In this study we have investigated the membrane behaviour of CCR5 expressed in fixed model cell lines using both structured illumination microscopy (SIM) and a new mode of imaging we have developed called Partially TIRF coupled HILO (PaTCH) microscopy. Using this combination of advanced biophysical techniques, we have been able to make observations into the clustering of membrane bound CCR5 as well as perform single-molecule investigations into the stoichiometry of these assemblies. Through the addition of perturbations employing the CCR5 agonist CCL5, we have been able to make preliminary observations into the ligand-dependent change in CCR5 behavior as proof-of-concept for the quantitative potential towards new biological insights with this approach.

### SIM investigations of CCR5

Initial investigations into the distribution of CCR5 foci were carried out using an established line of GFP-CCR5 expressing CHO cells that have been utilised in preceding studies. Our study aimed to unveil the distribution of CCR5 using super-resolution SIM microscopy, thereby allowing the precise localization of CCR5 for the analysis of its clustering behavior. The resulting 3D image data, which captured DyLight 650-MC-5 labeled CCR5 assemblies from the basal to the apical membrane, revealed that CCR5 collects into small puncta throughout the entire plasma membrane. Through the localization of the intensity centroids of these puncta we were able to quantify the level of clustering in the CCR5 distribution using Ripley’s H-function. Comparing these results with that of a randomly generated distribution of points, we found that CHO-CCR5 exhibits a clustered distribution with a modal clustering gradient of 0.004 ± 0.001. These results indicate that the puncta in which CCR5 appear to collect are in a non-random spatial distribution over the plasma membrane. Additional investigations are needed to determine whether the location of CCR5 puncta is correlated to a biological process and whether this organisation serves specific cellular roles. Finally, analysis of Ripley’s *H* values over a short range facilitated the determination of the nearest neighbor separation of CCR5 puncta with the mean distance being 290 ± 10 nm, a result that further guides the characterisation of CCR5 expression.

### PaTCH microscopy investigations of CCR5

To provide information on the structure of these CCR5 assemblies we performed single-molecule investigations using PaTCH microscopy. For this purpose, a new line of cells was created that stably expressed CCR5 at a level suitable for single-molecule microscopy. As well as low expression, this cell model required fluorescent labeling that could guarantee a ratio of one probe per CCR5 molecule for the purposes of molecular stoichiometry measurements. We therefore developed a line of CHO cells that stably express GFP-CCR5 and confirmed functionality and appropriate expression levels using flow cytometry-based calcium flux assays and immunolabeling. Utilising novel PaTCH microscopy to study this newly developed cell line, we were able to confirm the collection of CCR5 into small puncta and track the fluorescence intensity of these assemblies through time as they decayed due to photobleaching. Using this data and the method utilised in previous studies (Jin et al., 2021; Leake et al., 2006, 2008; Reyes-Lamothe et al., 2010; Syeda et al., 2019; Wollman et al., 2021, 2020b) we were able to determine the stoichiometries of individual CCR5 assemblies and form stoichiometry distributions for individual cell populations. We found that the distribution of CCR5 stoichiometries exhibits a significant range and can be characterised by the existence of periodic peaks in stoichiometry with an average interval of 2.3 ± 0.5 molecules. The existence of this periodicity leads us to believe that CCR5 puncta on the basal membrane likely consist of homodimeric sub-units, a result that is supported in the literature with other groups reporting the existence of dimeric CCR5 (Jin et al., 2018; Martínez-Muñoz et al., 2018b). Recent studies have been conducted that similarly employ CCR5 fused with GFP expressed within CHO cells, providing validity to this method of reporting (Li et al., 2021), however these studies employ GFP coupled on the C-terminus of CCR5, as distinct from the N-terminus coupling in our study. Due to the existence of a PDZ binding domain on the C-terminus of CCR5, C-terminal coupling raises potential concerns regarding the downstream effect of PDZ masking on the behavior of CCR5 (Delhaye et al., 2007; Hammad et al., 2010).

### Investigations of CCR5 after CCL5 interaction

Our investigations into the membrane behavior of CCR5 were extended through the perturbation of this model using the CCR5 agonist CCL5. This perturbation has been studied previously using bulk ensemble techniques which indicate progressive downmodulation of CCR5 (Signoret et al., 2005). Although our super-resolution SIM analysis did not detect a significant change in surface CCR5 following 5 minutes of ligand stimulation, we noted that the spread of stoichiometry values acquired through PaTCH analysis increased following stimulation, despite a similar level of overall tracked assemblies. The increase in frequency of both small and large stoichiometries, coupled with a decreased incidence of intermediate stoichiometries, could be associated with the movement of CCR5 subunits from their basal membrane location towards sites of internalisation. Through findings from previous investigations (Grove et al., 2014; Mueller et al., 2002; Signoret et al., 2004), these sites are suspected to be clathrin-coated pits in which CCR5 is theorised to report for the purposes of internalisation and recycling. Although a promising preliminary finding in our proof-of-concept study here, to provide further evidence for this model would sensibly require further single molecule investigation using varying levels of ligand perturbation. Through acquiring a dataset with varying perturbation times, one would aim to monitor further changes in the stoichiometry distribution coinciding with the downstream receptor internalisation seen in bulk ensemble investigations (see Supplementary Figure 2 b). Further, investigations including the fluorescent labeling of clathrin would provide insight into the colocalization of CCR5 and clathrin-coated pits.

### Summary

Through the application of complementary biophysical techniques, we have been able to perform super-resolved, single-molecule precise investigations of chemokine receptor CCR5 expressed in model cell lines. These investigations have provided hitherto unreported super-resolution images of CCR5 that allow us to reveal higher-order clustering of CCR5 subpopulations. Therefore, with the heterogeneity of GPCR subpopulations being linked with their signaling function (Martínez-Muñoz et al., 2018a), the ability of these techniques to characterise different structures and behaviors within independent sub-populations will facilitate the production of results relevant to the fields of both immunology and immuno-oncology. By utilising the suitability of these microscopy techniques for the investigation of live-cell imaging, we will gain the ability to track receptors as they travel throughout the plasma membrane, thereby quantifying the dynamic characteristics of CCR5 assemblies. In addition, our investigations can be readily augmented to include additional perturbations of both CCL5 and other CCR5 agonists and antagonists, thereby allowing single-molecule studies into the effects of CCR5 targeted drugs, such as Maraviroc. Finally, although the study of CCR5 within model cell lines allows the measurement of CCR5-ligand characteristics in the absence of competing binding partners, future extensions of our study will aim to employ the use of primary onco-immunogenic cells for the investigation of endogenously expressed CCR5, thereby enabling examination into the effects of a natural plasma membrane environment on the behavior of CCR5.

## Limitations of the study

In this study, we employed fixed model cell lines transfected to express CCR5. Although these model cells allow us to study CCR5 and its agonist CCL5 in the absence of competing binding partners, the biological relevance of this model system is limited by the fact that these are not primary cells. By extending this study to the investigation of CCR5 expressed endogenously in primary onco-immunogenic cells, we can perform a rigorous study that accounts for the presence of other chemokine receptors and ligands. Although this study aims to mitigate against effects induced by sample preparation, the extension of this study to investigate live cells would further mitigate any effect on the CCR5 distribution and cell morphology induced through formalin fixation and mounting and would allow for the determination of unhindered receptor dynamics. Emerging single-molecule optical microscopy techniques of single-molecule light sheet and single-molecule light field (Ponjavic et al., 2018) may prove valuable in the future to address questions of receptor dynamics and stoichiometry (Sims et al., 2020), by enabling potential visualisation of both apical and basal surfaces and 3D positions of single fluorophores to be determined from fast parallax measurements, respectively.

However, the extension of the methods discussed to the investigation of CCR5 within live tumor tissues introduces challenges associated with the imaging of optically thick samples, potentially comprising multiple cell layers, and the unavailability of stable receptor expression. This extension presents the opportunity for further development of the employed microscopy techniques, such as PaTCH, that are currently capable of imaging only a thin layer above the optical substrate, thereby allowing the investigation of single molecules above the basal membrane. Additionally, this study sees the employment of fluorescent labels that require exposure to high levels of laser excitation intensity, thereby raising potential concerns regarding the permanent photobleaching of reporters and phototoxicity effects to the cells. Although photodamage issues are mitigated in our study through the use of fixed end-point based experiments, any future extension to live cells will require the consideration of potential cell damage if cells are to remain viable in culture. Finally, despite the successful use of GFP and DyLight 650 dye to determine single-molecule, super-resolved stoichiometry and spatial localization of CCR5 in our study, these dye probes do possess room for improvement. Despite the enduring popularity of GFP, fluorescent proteins are comparable in size to CCR5 and have low fluorescence intensity when compared with modern organic dyes. When comparing the ligand-induced internalisation of GFP-CCR5 with that of CCR5, as investigated in previous studies (Signoret et al., 2004), we see a slower rate of internalisation that indicates a GFP-induced effect on receptor mobility. Through the introduction of SNAP-tag® or similar technologies into our cell model we may improve the overall signal-to-noise ratio of future acquisitions using small bright organic dyes that exhibit less perturbation to normal receptor function. Further, through the development of immunofluorescent labels that offer a dye to protein ratio of 1:1, we will be better equipped to perform measurements that report on the number of CCR5 molecules in distributions imaged using SIM.

## Acknowledgements

The authors thank members of the Leake and Signoret groups of the University of York for discussions and the Biosciences Technology Facility at York for assistance with flow cytometry. This work was funded by BBSRC (BB/T007222/1 and project 2279374) and EPSRC (EP/T002166/1). The authors also acknowledge funding from the UK’s Department for Business, Energy and Industrial Strategy through the National Measurement System.

## Data accessibility

Data associated with this study is freely accessible via DOI: 10.5281/zenodo.7082978

## Author contributions

Conceptualization: M.L, P.H, A.P-D, N.S

Project administration, Supervision and Funding Acquisition: M.L, N.S, M.S

Investigation, Methodology, Data curation, Analysis and Visualization: P.H, A.P-D, N.S, M.S

Software: A.P-D, P.H

Writing – original draft: P.H, A.P-D

Writing – review & editing: P.H, A.P-D, N.S, M.L, M.S

## Declaration of interests

The authors declare no competing interests.

## Methods

### Key resources table

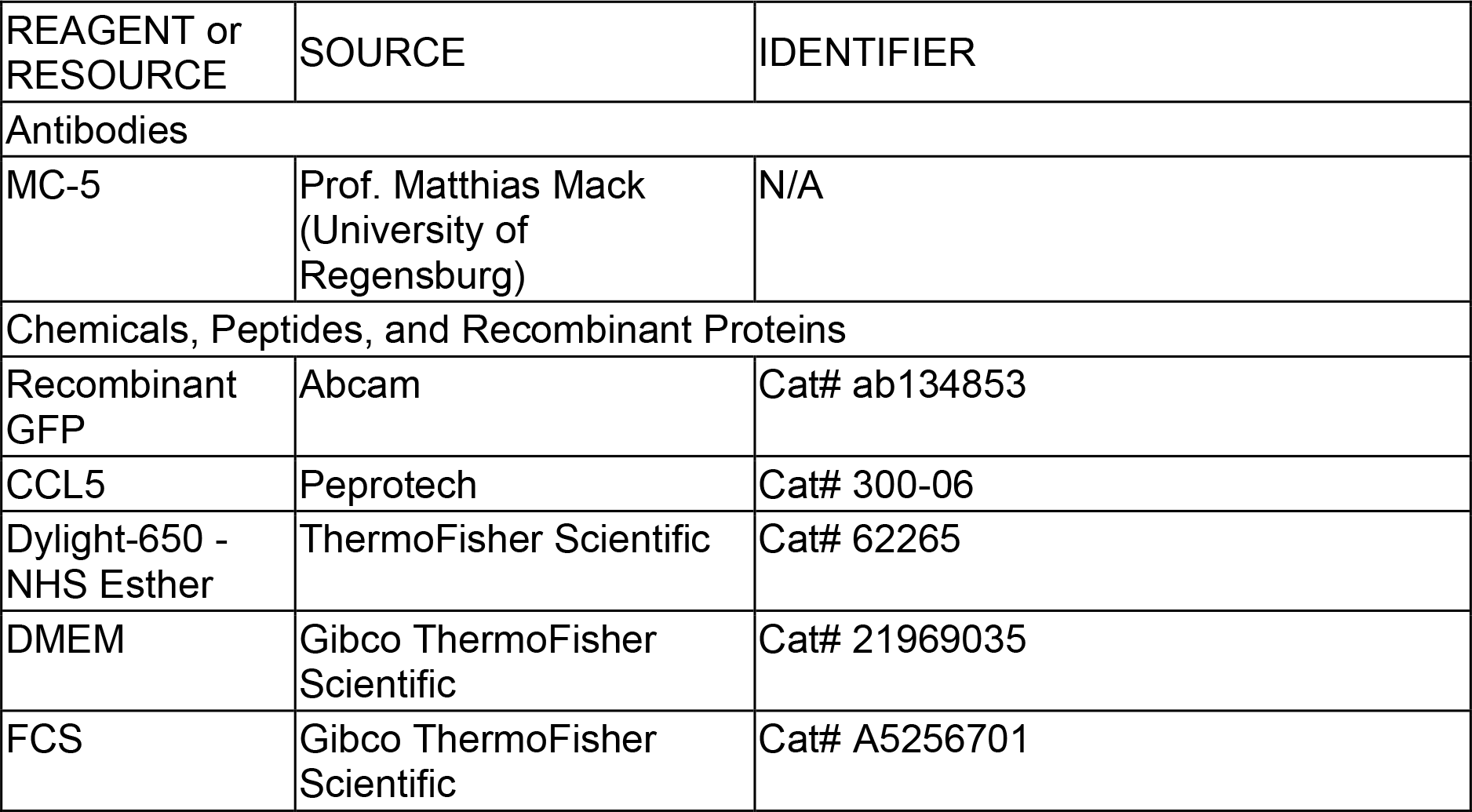
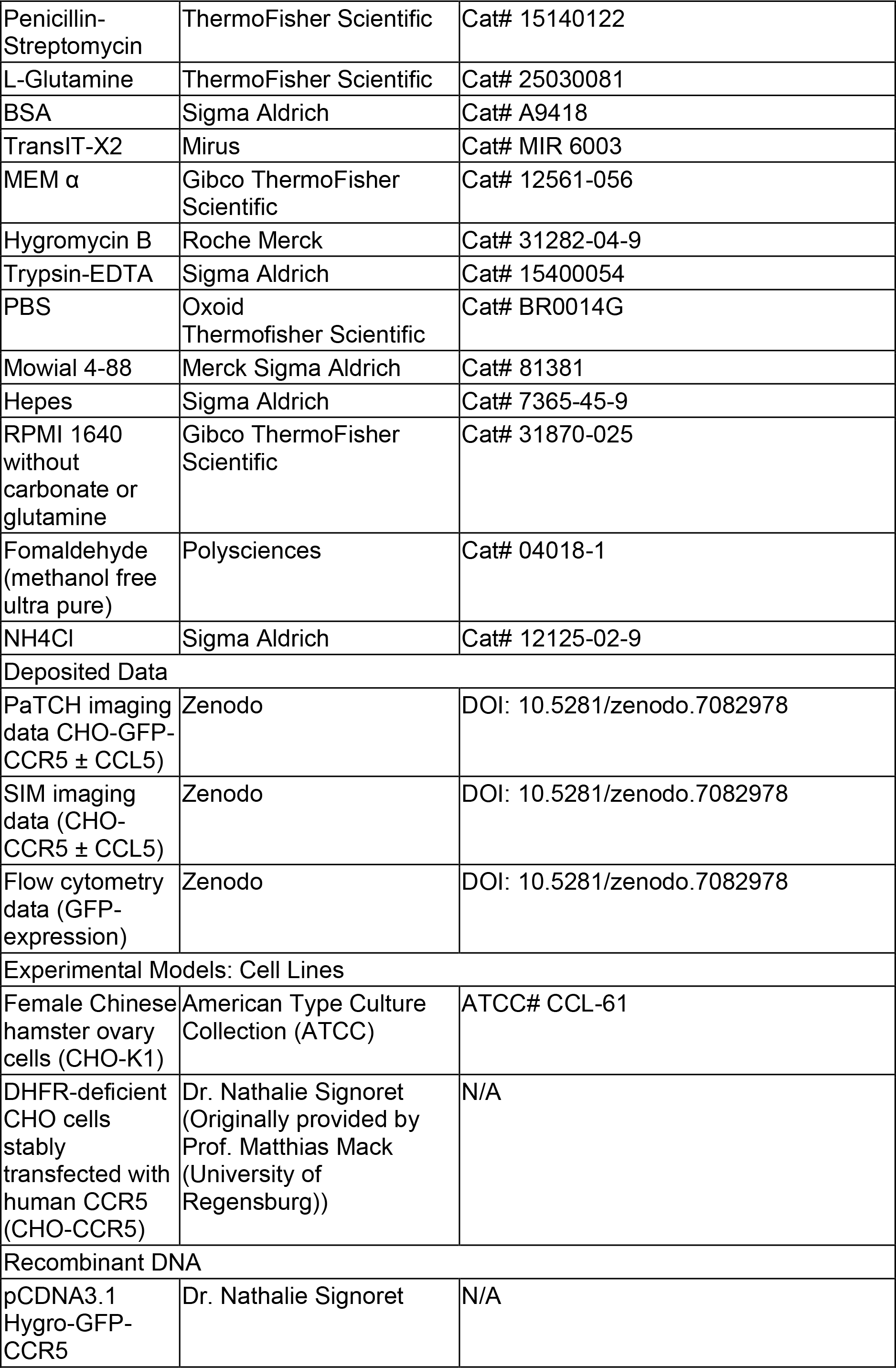
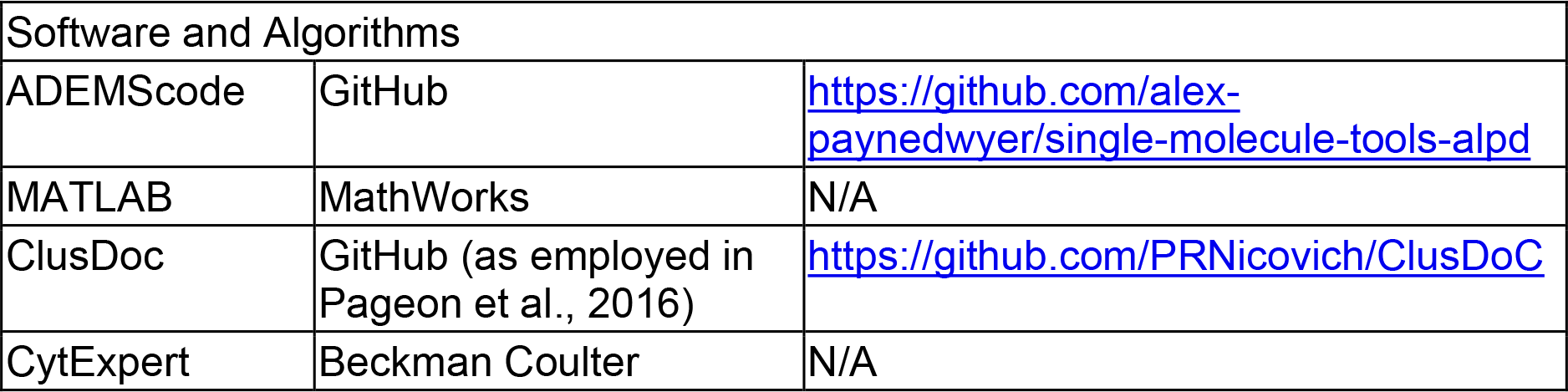

## Resource availability

### Lead contact

Further information and requests for resources and reagents should be directed to the lead contact, Mark C. Leake.

### Materials availability

This study did not generate new unique reagents.

### Data and code availability

- PaTCH imaging data, SIM imaging data and GFP-characterisation flow cytometry data have been deposited at Zenodo and are publicly available as of the date of publication. DOI is listed in the key resources table.
- All original code has been deposited at GitHub and is publicly available as of the date of publication. URL is listed in the key resources table.
- Any additional information required to reanalyse the data reported in this paper is available from the lead contact upon request.

## Experimental model and subject details

### Cell lines

Female Chinese hamster ovary cells (CHO-K1) and DHFR-deficient CHO cells stably transfected with human CCR5 (CHO-CCR5) (Signoret et al., 2005) were verified mycoplasma-free and maintained in complete (+ 10% FCS with 4mM L-glutamine, 100U/ml penicillin and 0.1mg/ml streptomycin) Dulbecco’s Modified Eagle Medium (DMEM) or MEM-alpha medium, respectively.

## Method details

### Materials

Tissue culture reagents and plastics were purchased from Invitrogen Life Technologies, (Paisley, UK), and chemicals from Sigma-Aldrich Company Ltd (Poole, UK), unless otherwise indicated. CCL5 (RANTES) was purchased from PeproTech EC Ltd.

### Transfection and generation of new cell line

CHO-GFP-CCR5 cell lines were generated by transfecting CHO-K1 with a pCDNA3.1 Hygro-GFP-CCR5 (Gómez-Moutón et al., 2004) construct in the absence of a strong pre-promoter amplification region, using the TransIT-X2 transfection reagent from Mirus (MIR 6003). CCR5 expression was maintained by culturing cells in the presence of 400 μg/ml hygromycin and GFP-CCR5 expressing cells were isolated for single-cell cloning through the process of serial dilution cloning. In this process, single GFP-CCR5 positive cells were seeded individually in a 96-well plate and expanded by continuous culture in the presence of hygromycin. Selection of specific populations based on their clonality and relatively low GFP signal intensity was aided through the characterisation of GFP expression using Flow cytometry (CytoFLEX LX, Beckman Coulter) against non-transfected CHO-K1 controls, as shown in Figure 3. Analysis of Flow cytometry results was performed using CytExpert (Beckman Coulter).

### GFP-CCR5 CHO cell preparation for PaTCH microscopy

GFP-CCR5 CHO cells from an 80% confluent well of a 6-well plate were detached in Trypsin-EDTA and seeded on 1.5 mm thick round coverslips at a dilution of 1:10 3 days prior to mounting. Cell samples were fixed with a solution of 3% formaldehyde in Phosphate Buffer Saline (PBS) for 20 mins at room temperature, before being extensively washed in PBS. Coverslips were mounted in Mowiol, as previously described in preceding studies (Kasprowicz et al., 2018).

### Immunofluorescence staining of CHO-CCR5 cells for SIM

CHO-CCR5 cells from an 80% confluent 10 cm dish detached in Trypsin-EDTA were seeded on 1.5 mm thick round coverslips two days prior to mounting. The culture medium was then replaced with Binding Medium (BM: RPMI 1640 without carbonate or glutamine, 0.2% (w/v) BSA, 10 mM HEPES adjusted to pH 7). For chemokine treatment, CCL5 was added to the BM before incubating coverslips at 37°C for 5 min at a saturating concentration of 100 nM (Andrews et al., 2008; Combadiere et al., 1996; Mack et al., 1998; Proudfoot et al., 2003; Signoret et al., 2005). Incubation was stopped by fixing samples as described above. For staining, free aldehyde groups were quenched with a 50 mM NH4Cl solution for 20 mins before saturation of non-specific binding in PBS with 1% FCS (PBS/FCS). Cells on coverslips were labeled intact with DyLight 650-MC-5 (2 μg/ml) in PBS/FCS for an hour before washing in PBS and mounting samples in Mowiol, as described above. MC-5 is an anti-human CCR5 mAb grown from a hybridoma and purified by Prof. Matthias Mack (Signoret et al., 2000). MC-5 non-interference with CCL5 binding, CCR5 conformation, activation and internalisation has been validated by numerous studies (Blanpain et al., 2002; Signoret et al., 2005, 2000) and has been previously used to follow chemokine-mediated CCR5 stimulation using TIRF (Grove et al., 2014). MC-5 was fluorescently labeled using DyLight 650 NHS ester coupling kit (Thermo Fisher) with a dye:protein coupling ratio of 1.57:1.

### PaTCH imaging

The Slimfield microscope (without alteration for PaTCH acquisition) is based on a custom-built epifluorescence/TIRF optical pathway previously described (Payne-Dwyer and Leake, 2022; Plank et al., 2009; Syeda et al., 2019). An optimised angle of excitation beam delivery distinguishes PaTCH imaging from the traditionally used TIRF and HILO microscope settings, thereby facilitating the imaging of transmembrane proteins in mammalian cells (Wollman et al., 2022). In this study the angle of incidence of the excitation beam was set to a sub-critical angle of 55° by translation of a telescope lens, as calibrated using the lateral displacement of the beam downstream of the focal plane (Dresser et al., 2021). At this angle, we estimated that ~30% of the incident light is coupled into a reflected TIRF mode, with the associated enhancement of the excitation field at the surface. The remaining light couples into transmitted HILO modes, which extend the excitation field into regions of the basal membrane not directly contacting the coverslip, but not the interior of the cell.

The 488 nm wavelength laser (Coherent OBIS LX) was spatially filtered to the TEM_00_ mode and delivered by an oil-immersion objective lens (Nikon ApoTIRF, 100×, NA 1.49). The illumination covered an area c. 60 μm wide (diameter at 1/e^2^ peak intensity) in the sample plane as characterised using a sample of immobilised standard fluorescent microbeads (Promega). The source power was 30 mW, corresponding to an excitation intensity of approximately 0.5 kW/cm^2^ at the sample.

A single image sequence was captured for each field of view in OME TIFF format using a Prime 95b camera (Teledyne Photometrics). The exposure time was 10 ms per frame, during which the laser was digitally triggered, and a total framerate of 77 fps over 1000 - 3000 frames at 53 nm/pixel magnification. The estimated lateral resolution is 180 nm, while the localization precision of each focus is approximately 40 nm (Lenn and Leake, 2012). The detection performance was characterised using *in vitro* recombinant GFP immobilised to a coverslip (Delalez et al., 2010; Leake et al., 2006; Wollman et al., 2020a).

### Single particle tracking

The custom software suite, ADEMScode (MATLAB, MathWorks) (Miller et al., 2015) was employed to detect local maxima inside circles of 8-pixel radius (foci), above background (averaged over 17 pixel squares) in each frame of a given image sequence. These foci were then thresholded using signal-to-noise ratio and were fitted with a Gaussian intensity mask to establish the super-resolved centroid, width and integrated intensity (background subtracted sum of pixel values). The foci were then linked into tracks based on their persistent overlap (75-100%) and intensity ratio (50-200%) across adjacent frames. A representative example of these super-resolved tracks overlaid on the parent PaTCH image is shown in Supplementary Figure 3 a. The initial intensity of each track was linearly extrapolated back across the first five frames to the time point of initial laser exposure. This initial intensity was divided by the characteristic integrated intensity of a focus containing a single GFP in order to determine the number of GFP molecules present in each track, which, given the 1:1 labeling of the protein of interest, was equated with the CCR5 stoichiometry. Only tracks determined in the first 10 frames of laser exposure were then used for estimates of stoichiometry to avoid undercounting due to photobleaching. The characteristic brightness of a single GFP was estimated from the modal brightness of GFP-CCR5 foci after sufficient photobleaching, to ensure the presence of single GFP molecules. This value of brightness was confirmed to be broadly consistent with values gained from recombinant GFP immobilised to a coverslip as well as estimates of the GFP brightness determined from the monomeric intervals in intensity due to stepwise photobleaching of GFP-CCR5 foci, as identified by a Chung–Kennedy edge-preserving filter (Leake et al., 2004, 2003; Payne-Dwyer et al., 2022; Wollman and Leake, 2015). Representative examples of single molecule intensity traces towards the end of the photobleaching process are shown in Supplementary Figure 3 b, with an inset trace demonstrating the accuracy of the estimation of the brightness of a single GFP probe. The collated stoichiometry of all tracks were represented as kernel density estimates with kernel width of 0.6 molecules, corresponding to the root mean squared (rms) detection sensitivity of the integrated intensity of a single GFP molecule in a focus (Figure 5). The periodicity was calculated by sorting the stoichiometries of a population of tracks (Wollman et al., 2017) and taking the nearest-neighbour differences (Payne-Dwyer et al., 2022). These intervals were themselves then plotted as a kernel density estimate with kernel width of 0.6 molecules. The periodicity was quoted as the modal peak in this distribution (Figure 5 inset), with the error estimated as 0.6 molecules, multiplied by the square root of the ratio of the mean stoichiometry and the number of extrapolation points, divided by the number of tracks under the main peak.

### SIM imaging

Super-resolution imaging was performed using a custom SIM system built around an inverted widefield epifluorescence microscope (IX71, Olympus) as described in (O’Holleran and Shaw, 2014; Shaw et al., 2015). Illumination patterns were generated by projecting the spatially filtered image of a binary phase grating, displayed on a liquid-crystal-on-silicon spatial light modulator (SLM) (SXGA-3DM, Forth Dimension Displays), into the sample. Images were recorded using a scientific CMOS camera (Flash 4.0, Hamamatsu Photonics), with the global exposure period of the camera’s rolling shutter synchronised to the SLM. Each super-resolution image was reconstructed from nine raw images captured under illumination of the sample with a series of sinusoidal excitation patterns (three pattern orientations separated by 120° and pattern phases separated by 2*π*/3 per orientation). High pass filtering was applied to suppress out-of-focus information close to the centre of each separated Fourier space information passband (O’Holleran and Shaw, 2014) before passbands were shifted and combined as described in (Gustafsson et al., 2008). Images of Dylight 650-MC-5 labeled CHO-CCR5 were acquired using a quad band fluorescence filter cube (TRF89901-EM, Chroma) and a 60×/1.3 silicone immersion objective lens (UPLSAPO, Olympus) with excitation at 638 nm using a diode laser (Luxx, Omicron), with an effective lateral (Abbe) resolution of approximately 170 nm. Post reconstruction, images captured at different focal offsets were corrected for photobleaching by scaling using an exponential decay curve measured by repeated imaging of the same region of a cell under identical conditions.

### Clustering analysis

Images of CHO-CCR5 labeled with DyLight 650-MC-5 obtained using SIM underwent cell boundary segmentation to remove extracellular foci. Images then underwent binarization using a combination (in parallel, followed by an AND operation) of global Otsu thresholding and local Otsu thresholding with a rolling ball radius of 25 pixels. The binary images revealed objects corresponding to local enrichment of CCR5. The number of objects, and the characteristic properties of each object, including area, centroid and circularity were determined using the *Analyze Objects* function in ImageJ/Fiji. The number density of objects (Figure 2 d) was calculated using a 2D kernel density estimate of the object centroid coordinates, with kernel width of the widefield lateral resolution, ~180 nm.

Centroid coordinates of foci were analysed using ClusDoc software (Pageon et al., 2016). The relative clustering or dispersal of objects was assessed for each individual cell using the Ripley H function. The Ripley H function is equal to the Ripley L function less the radius, H(r)=L(r)-r, where L(r) is the radius of a circle in which the experimentally counted points inside would otherwise be uniformly distributed (Kiskowski et al., 2009). Thus, Ripley H is a measure of the deviation from uniform distribution, H(r)=0, with positive deviations corresponding to ‘attractive’ clustering of objects. The clustering results were validated against simulated negative and positive controls, which were point clouds generated respectively from random 2D coordinates or points on a square lattice with rms noise of half the lattice spacing (MATLAB). Nearest neighbour distances were calculated using the minimum value of H(r) associated with the initial negative H(r) values, values were averaged over all cells to provide a mean value ± standard error of the mean. Clustering gradients were determined as the peak of a kernel density estimate of gradient values calculated pairwise between adjacent H(r), with a kernel width of 0.001.

### Quantification and statistical analysis

MATLAB was used for statistical tests as reported in the Results section. To account for multiple comparisons (across ≤ 5 tests per sample: intensity/stoichiometry, periodicity and cluster density), we used a conservative Bonferroni-adjusted significance level of ɑ = 0.05/5 = 0.01. Nonparametric statistical tests were used to test for significance, chiefly the Brunner-Menzel test with exact values of *N* and *p* reported. *N* represents in each case either the number of foci or the number of cells.

As the underlying distributions and variances were unknown *a priori*, a target sample size of n ≥ 5 cells (and n ≥ 24 tracks) per condition was estimated. These targets were based on a minimum detection level of 1 s.d. at 80% power under normal statistics for a one-tailed or two-tailed Z-test respectively. Cell cultures were assigned randomly for ligand +/− groups. Of 24 total SIM (21 PaTCH) acquisitions, 23 (21) were included in analysis based on sufficient quality of the initial microscope focus. The lower target number of cells reflects the experimental throughput of the microscopy techniques.

## Notes

### Competing Interest Statement

The authors have declared no competing interest.

### Summary of Updates

Additional experiments reports; additional analysis performed; additional references added; text reworded

