## Supplementary Information for "Single-molecule and super-resolved imaging deciphers membrane behaviour of onco-immunogenic CCR5"

### Super-resolved imaging deciphers ligand dependent membrane behaviour of the onco-immunogenic CCR5 receptor.

### Supplementary movie legends

***Supplementary Movie 1. A 3D movie of a CHO-CCR5 cell shows distinct puncta throughout the membrane.*** *3D reconstruction of 7 Individual SIM images of a CHO-CCR5 cell labeled with DyLight 650-MC-5, as shown in Figure 1 a-g). (Scale bar 2 µm).*

***Supplementary Movie 2. A 3D movie of a CHO-CCR5 cell after ligation shows retention of distinct puncta throughout cell.*** *3D reconstruction of 7 Individual SIM images of a CHO-CCR5 cell labeled with DyLight 650-MC-5 and perturbed with CCL5, as shown in Figure 6 a-g). (Scale bar 2 µm).*

### Supplementary figures

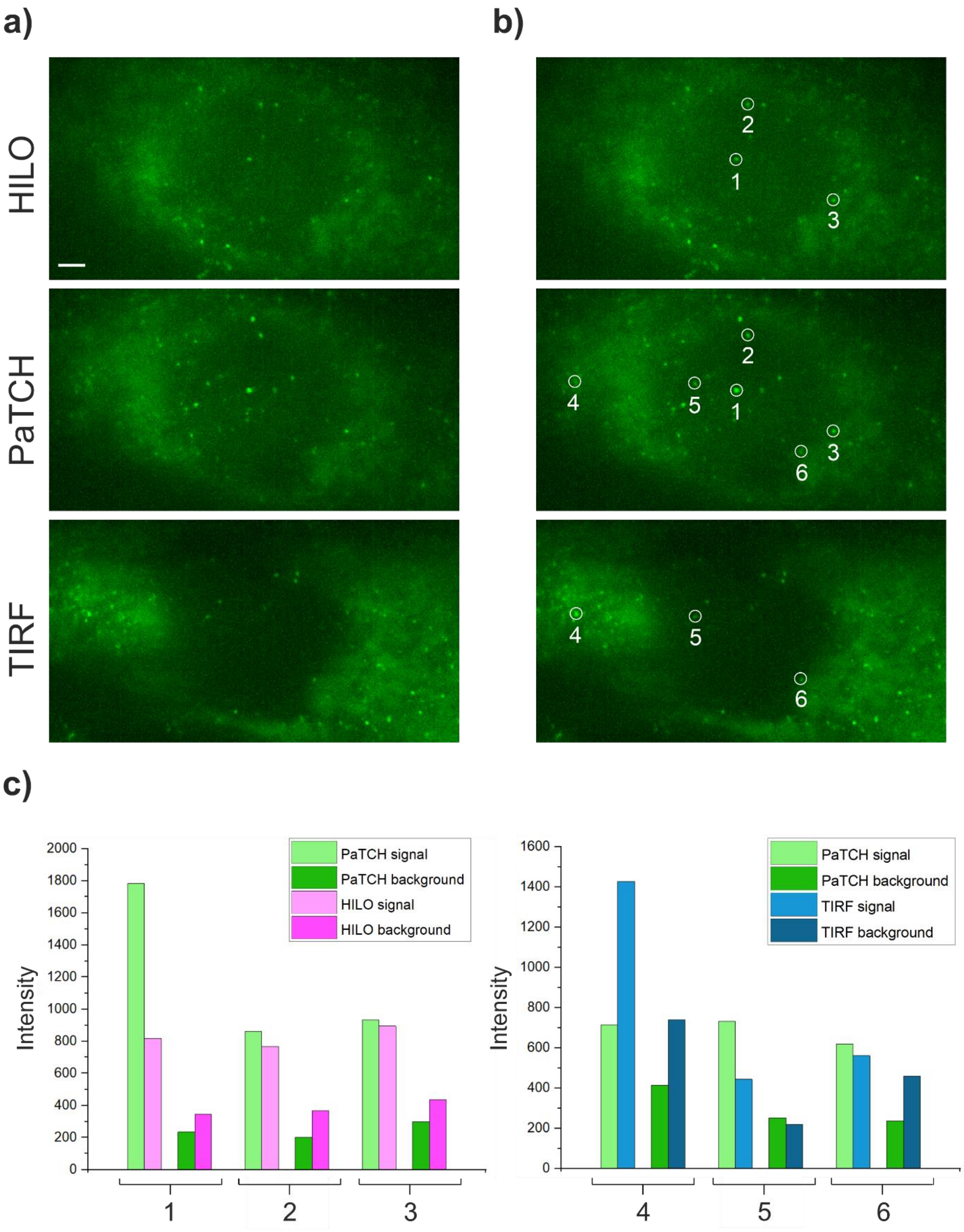


***Supplementary Figure 1. Comparison of HILO, PaTCH and TIRF imaging modes for the single molecule detection of GFP-CCR5 in CHO-GFP-CCR5.*** *a) GFP-CCR5 expressing CHO cell imaged using HILO, PaTCH and TIRF microscopy. Thereby revealing the increased signal of puncta within the basal membrane in PaTCH when*

compared with HILO, whilst demonstrating the uniformity of excitation of the basal membrane in PaTCH when compared with TIRF. (Scale bar 2 µm). b) Circular overlays highlighting CCR5 assemblies present in both HILO/PaTCH images and in both TIRF/PaTCH images. Overlays are numbered to facilitate further analysis. c) Comparison of signal and background between HILO and PaTCH imaging modes, in puncta labelled 1-3, and between TIRF and PaTCH imaging modes, in puncta labelled 4-6. Intensity represents the raw integrated density, captured using a 6-pixel diameter circle, above a mean global background calculated using the extracellular space. Measurements of signal were taken directly over the puncta, while measurements of local background were taken adjacent to puncta. In general, puncta imaged using PaTCH benefit from a signal enhancement, relative to HILO, due to the TIRF-coupled component of illumination. Further, although TIRF is capable of providing enhanced signal relative to PaTCH, as seen in puncta 4, this restricted illumination mode results in higher background from fluorescent material close to the coverslip as well as lower signal from puncta not in close contact with the coverslip. These results combined with the result for the average signal to background ratio of 1.8 ± 0.1, 3.6 ± 0.6 and 2.4 ± 0.4 for puncta within HILO, PaTCH and TIRF respectively demonstrate the general increase in signal and reduction in background found in PaTCH when compared with HILO and TIRF.

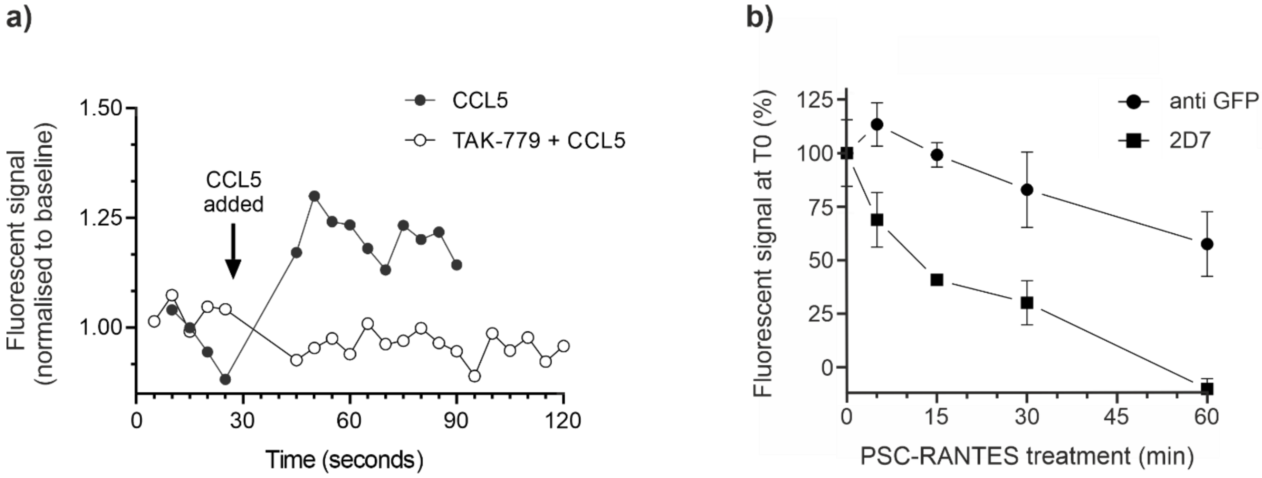


***Supplementary Figure 2. Increase in calcium flux, coupled with a decrease in anti GFP and 2D7 antibody binding, upon ligand stimulation confirms the functionality of GFP-CCR5.*** *a) Calcium flux assay in which the change in calcium associated fluorescent signal is monitored within samples of CHO-GFP-CCR5 during live exposure to 10 nM CCL5, with and without pre-exposure to the CCR5 antagonist TAK-779. Thereby revealing an increase in CCL5-associated calcium signalling in the absence of an antagonist, suggesting a functional response of GFP-CCR5 to CCL5. Values of fluorescent signal are reported normalised to the average baseline fluorescence prior to CCL5 exposure. b) Fluorescent signals associated with antibodies bound to GFP (anti GFP) and the CCR5 chemokine binding site (2D7) are measured within CHO-GFP-CCR5 cells that underwent fixation after varying levels of*

*exposure to the super-agonist PSC-RANTES at a concentration of 100 nM. Thereby revealing a decrease in both the accessibility of the GFP epitope and the availability of the chemokine binding site, suggesting the binding of ligand and the subsequent internalisation of GFP-CCR5.*

**a)**

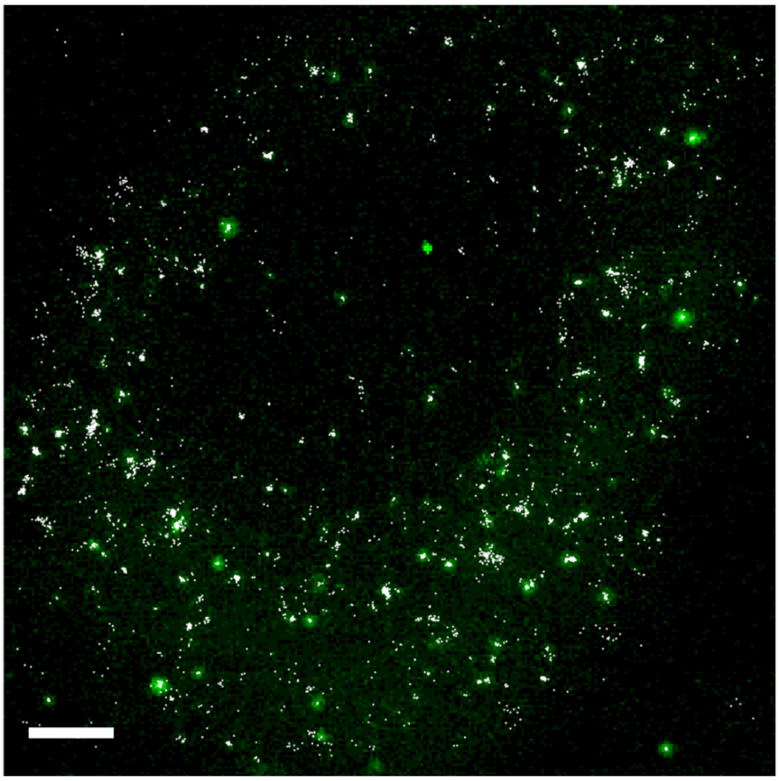

**b)**

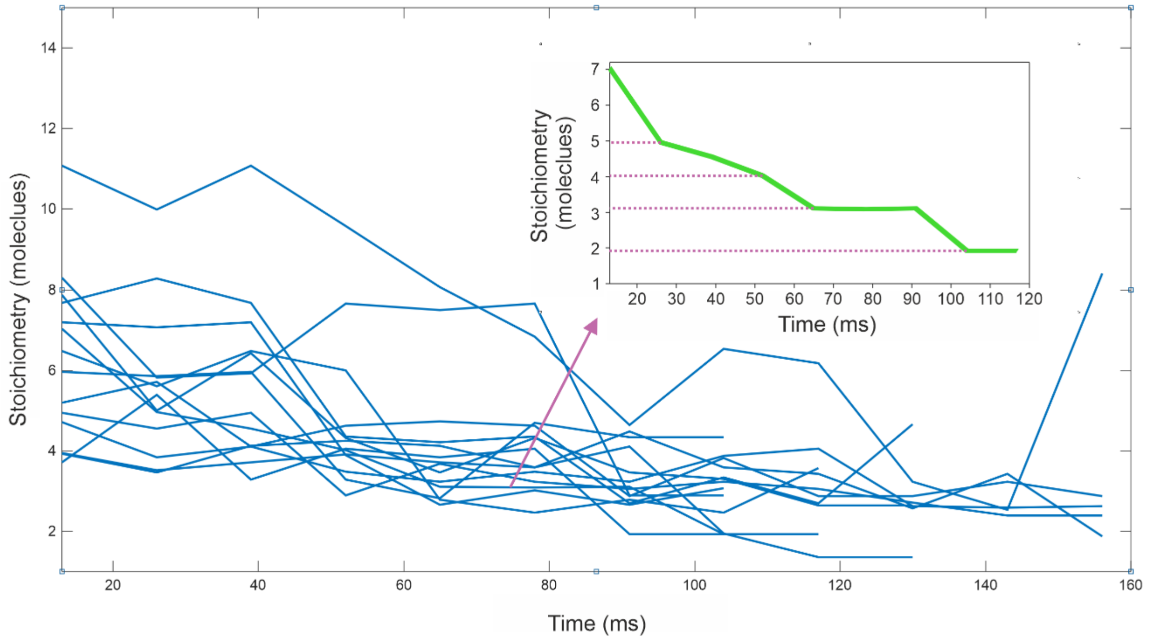


***Supplementary Figure 3. ADEMScode is used to detect foci within PaTCH microscopy images.*** *a) GFP-CCR5 expressing CHO cell imaged using PaTCH*

*microscopy with an overlay (white) showing tracks determined by ADEMScode tracking (MATLAB). (Scale bar 2 µm). b) Chung–Kennedy edge-preserving filtered Intensity time traces revealing the photobleaching-induced intensity decay of tracked foci towards the end of the photobleaching process. The representative traces shown here exhibit fluctuations in intensity, however this effect is accounted for in the determination of stoichiometry. Inset trace (green) shows an example of a focus whose intensity underwent decay with minimal fluctuation, dropping in a stepwise fashion from an apparent stoichiometry of 7 to 2, thereby supporting the accuracy of the estimated brightness of a single GFP molecule acquired from the modal brightness of monomeric GFP-CCR5.*

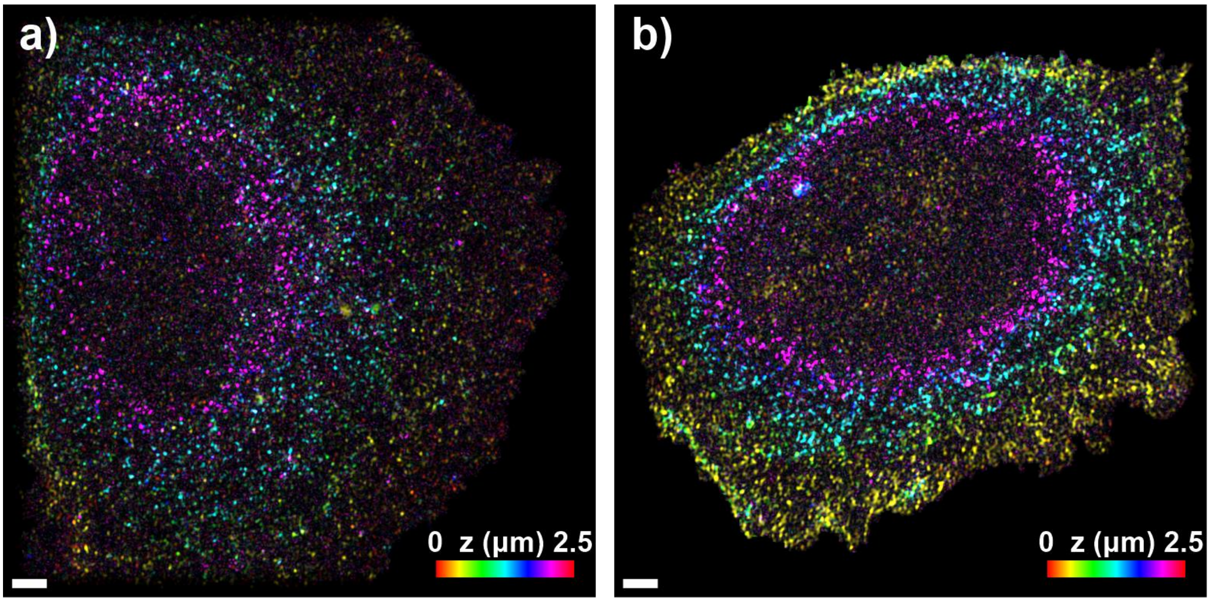


***Supplementary Figure 4. Comparison of cellular imaging demonstrates slight variation in brightness.*** *Color depth projection of cell images shown in a) Figure 1 h) and b) Figure 6 h). Comparison of which at identical contrast settings demonstrates the slight variation in brightness that can exist between cells. This change in brightness can stem from many factors, including the natural variance of expression in this non-clonal cell model. (Scale bar 2 µm).*

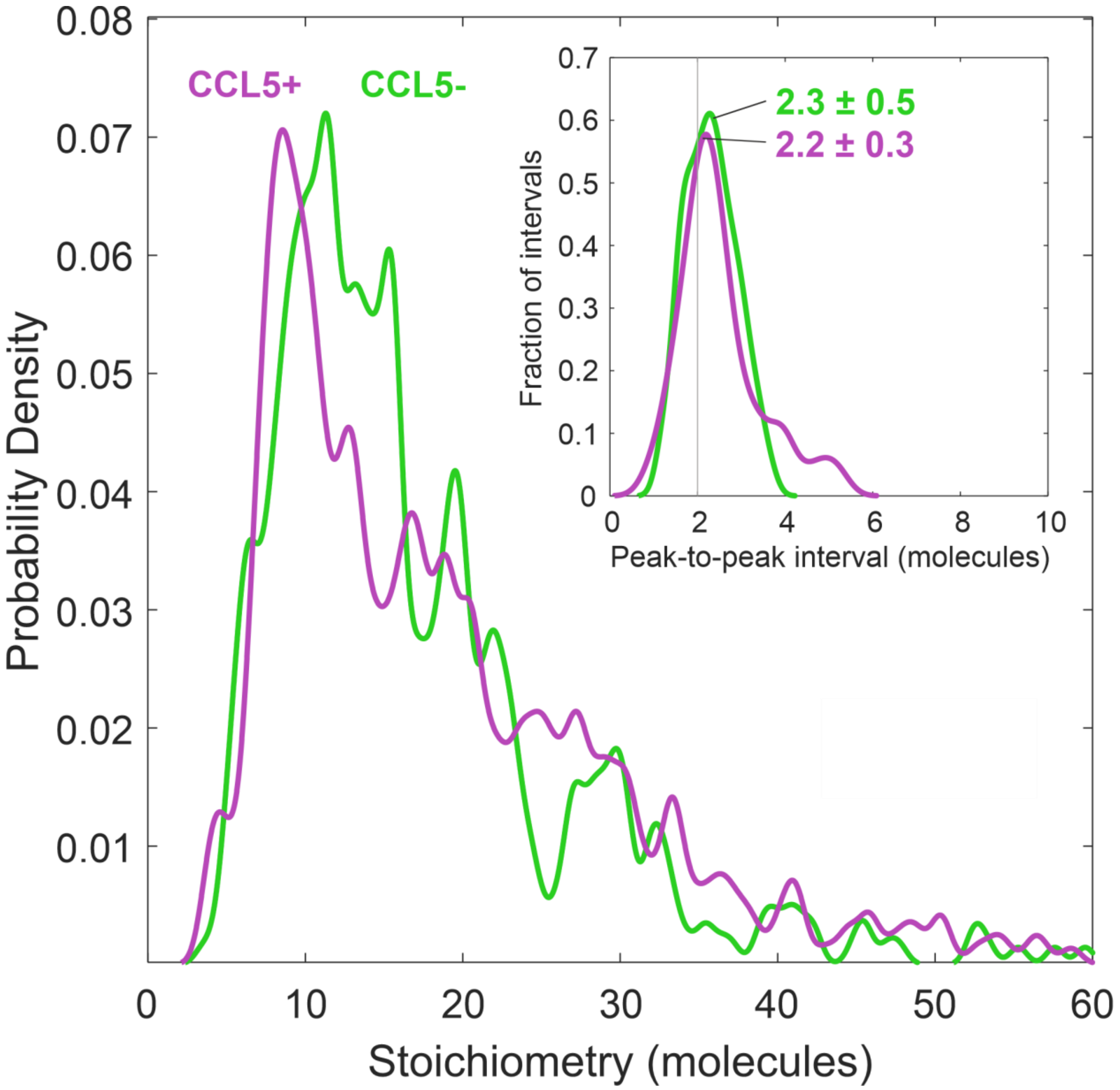


***Supplementary Figure 5. Periodic stoichiometry distribution indicates that CCR5 assemblies comprise dimeric subunits both before and after CCL5 exposure**. Kernel density estimates of stoichiometry and (inset) periodic stoichiometry intervals of GFP-CCR5 associated foci before (green) and after (magenta) the addition of CCL5 (N=460 or 507 tracks respectively) detected by PaTCH microscopy in GFP-CCR5 transfected CHO cells (N=9 and 11 cells respectively). Kernel width = 0.6 molecules, corresponding to the total uncertainty in the single molecule stoichiometry, rather than statistical fluctuations. Measured intervals in probability density or stoichiometry are therefore more reliable at lower stoichiometry. (BM test for difference in periodicity, p=0.479* | *NS).*

### Supplementary methods

<u>Comparison of HILO, PaTCH and TIRF microscopy</u>

A slimfield microscopy experiment was performed on fixed samples of CHO-GFP-CCR5 using a varied angle of incidence of the excitation beam (see Supplementary Figure 1). This experiment was performed under the same conditions as the PaTCH experiments outlined in the methods section. Individual cells first underwent an epifluorescent prebleach lasting 750 frames (~10 s) to minimize the effect of photobleaching on the comparison of imaging modes. These cells were then imaged for 100 frames using angles of incidence of 45°, 55° and 62° sequentially, corresponding to HILO, PaTCH and TIRF respectively. Acquired stacks of images were used to form maximum z intensity projections of singular cells under all imaging modes. The location of puncta within projected images was determined using intensity thresholding, allowing the spatial distributions of puncta within PaTCH images to be transformed onto HILO and TIRF images, thereby facilitating the determination of puncta present across multiple imaging modes. The signal of isolated puncta and the adjacent background were determined under a circular window of 6 pixels in diameter. Intensity of signal and background is represented as the raw integrated density, captured under the circular window, normalized to the global background of the sample. Global background was calculated as the average raw integrated density in the extracellular volume across the three imaging modes. Finally, average values of the signal to background ratio were calculated using 10 puncta from each of the three imaging modes in order to provide a mean value ± standard error of the mean.

<u>Characterization of new cell line using calcium flux assay:</u>

Calcium flux assays were performed to confirm the functionality of CCR5 within the CHO-GFP-CCR5 cell line (see Supplementary Figure 2 a). Cells were detached in PBS 10mM EDTA, washed twice in PBS without calcium before resuspension in PBS at 2 x $10^6$ cells/ml. Quest Fluo-8 AM was added to give a 2.5 μM concentration and cells were incubated for 30 minutes in the dark at room temperature. Excess dye was removed by two PBS washes before cell resuspension at 1 x $10^6$ cells in HBSS medium containing 1.26 mM calcium chloride. Intracellular calcium concentration changes within 500 μl aliquots of cells in response to 10 nM CCL5, the known optimal concentration for calcium flux assays(Combadiere et al., 1996), were determined by analysis of cell fluorescence on a Cytoflex S flow cytometer (Beckman Coulter) using an argon laser at a wavelength of 488 nm. Labelled cells were kept on ice as 500μl aliquots until use, some of which were pre-exposed to 400 nM of the CCR5 antagonist TAK-779. Cell aliquots were aspirated before stimulus was added in order to define the baseline fluorescence of the sample. Acquisition was resumed for the duration of the response and repeated, as required. A blank stimulation was used to control for the mechanical impact of instrument on readings. Collected data were analyzed by plotting the fluo-8-$Ca^{2+}$- FITC fluorescence signal against time, with successive gates of 5 second intervals for the duration of the response. Results are reported as fluorescence normalized to the baseline signal of each sample before stimulation and graphs were plotted in Prism v9.4.1 (GraphPad Software Inc., La Jolla, USA).

<u>Confirmation of GFP-CCR5 internalisation at extended ligand exposure:</u>

A flow cytometry-based assay was performed to confirm the internalization of GFP-CCR5 after extended exposure to super-agonist PSC-RANTES (gift from O. Hartley, University of Geneva) (see Supplementary Figure 2 b). Assay was performed in Binding medium [BM: RPMI 1640 without bicarbonate, containing 0.2% BSA and 10 mM Hepes, pH 7.0]. Cells were detached in PBS 10mM EDTA and resuspended at $2 \times 10^6$ cells/mL and incubated in BM alone or containing 100 nM PSC-RANTES for up to 1 hour at 37°C. A 100 µl aliquot of the cell suspension was taken for each time point and transferred to a 96W plate kept on ice. At the end of the time course, each sample was split between two wells for antibody labelling with 2D7, a CCR5-specific monoclonal antibody recognising the chemokine binding site, and with an anti-GFP monoclonal antibody in FACS Buffer [PBS/1%FCS/Azide]. Samples were fixed overnight in FACS buffer plus 1% formaldehyde. Bound antibodies were then detected using GAM-Dylight 650 in FACS Buffer, and all samples were analysed using a CytoFlex S flow cytometer (Beckman Coulter). For each antibody staining, the percentage of fluorescent signal was calculated from the specific mean fluorescence intensity (MFI= CCR5 antibody MFI subtracted for background).
